# Geometry-preserving vector field reconstruction of high-dimensional cell-state dynamics using ddHodge

**DOI:** 10.1101/2025.04.16.649050

**Authors:** Kazumitsu Maehara, Yasuyuki Ohkawa

## Abstract

The differentiation potency of cells is governed by dynamic changes in gene expression, which can be inferred from single-cell RNA sequencing (scRNA-seq) data. While velocity-based approaches have been used to analyze cell state changes as vector fields, extracting acceleration (change of change) information remains challenging because of the sparsity and high dimensionality of the data. Here, we developed ddHodge, a framework based on Hodge decomposition for precise vector field reconstruction. ddHodge accurately recovers all basic components of the vector field, namely, the gradient, curl, and divergence, including the acceleration of the cell state, as second-order derivatives, even from biased and sparse samples. Furthermore, we extend the method to approximate high-dimensional gene expression dynamics on lower-dimensional data manifolds. By applying ddHodge to scRNA-seq data from mouse embryogenesis, we revealed that the gene expression dynamics during development follow a gradient system shaped by potential landscapes, which has not previously been validated with real data. Furthermore, we quantified differentiation potency as cell state stability on the basis of the divergence and identified key genes that drive potency. Our general computational framework for analyzing complex biological systems can elucidate cell fate decisions in developmental processes.

## Introduction

The dynamics of cell states, namely, the temporal changes in cell states, underpin all biological phenomena, including development, regeneration, differentiation, and pathology. One of the primary goals of life sciences is to understand these cell state dynamics. However, these dynamics represent a high-dimensional, multibody system involving interactions among tens of thousands of proteins, nucleic acids, and lipid molecules, constituting a complex, nonequilibrium system^1,2^. Consequently, traditional hypothesis-driven approaches have focused on small subsystems formed by the interactions among the molecules of interest, creating building blocks for understanding the whole system. Recent advancements in single-cell omics^3–6^ have enabled comprehensive evaluations of cell states represented by all gene expression profiles, which could allow a data-driven understanding of complex cell state dynamics. However, single-cell measurement techniques are invasive, and the resulting data are static, lacking temporal changes for each cell. Therefore, pseudotime analysis techniques^7–9^ have been employed to construct temporal information based on of large-scale static snapshot data to analyze gene expression dynamics. Nevertheless, conventional pseudotime analysis methods provide only an order of time, and the speed and acceleration/deceleration cannot be evaluated, making it challenging to analyze detailed dynamics, including the stability of states that represent sensitivity of gene expression responses to perturbations, as well as more diverse types of state changes.

Therefore, RNA velocity^10,11^ has frequently been used to trace changes in cell states as trajectories in gene expression space by utilizing the time derivative of the gene expression (velocity) of each cell. Based on this time-derivative inference approach involving gene expression data, methods for reverse engineering these dynamics (vector field reconstruction) have been proposed, allowing analyses of the interactions among genes that drive local dynamics and the least action paths of the cell state^12^. However, owing to the lack of a sufficient theoretical foundation to ensure accurate reconstruction of vector fields on the basis of high-dimensional data, a methodology for precisely reconstructing cell state dynamics, appropriately summarizing the dynamics occurring in high-dimensional space, and evaluating the stability of each cell state has yet to be developed.

Here, we developed a high-dimensional dynamics analysis framework called ddHodge. Our framework is based on the mathematical theorem of Hodge decomposition, which provides the three fundamental components of vector fields on manifolds^13,14^. ddHodge offers a unified perspective on the problem of high-dimensional vector field reconstruction on manifold and provides a practical method to precisely reconstruct dynamics in unprecedented detail. Through simulations and validation using single-cell RNA sequencing (scRNA-seq) data, we demonstrated that ddHodge can provide a summary of the overall unknown high-dimensional dynamics and identify genes associated with fate determination by affecting the stability of cell states.

## Results

### ddHodge provides a framework for analyzing the high-dimensional dynamics of cell states

The cell states represented by each data point are generated from an unknown differential equation system ***F***(x, y) (Figure 1A, left). These states are pairs of observed positions (x, y) and their corresponding velocities ***F***(x, y). Each velocity is given as the gradient of an unknown potential function U. The magnitude of the potential value at each point reflects the temporal ordering of the cell states (Figure 1A, middle). The convex region in the potential landscape, indicates the stability of states that are diversifying; branching is likely to occur, and the concave region is considered to be converged; the cell states adopt specific lineages (Figure 1A, right). Therefore, the shape of the potential landscape is a critical indicator of dynamics.

**Figure 1.**
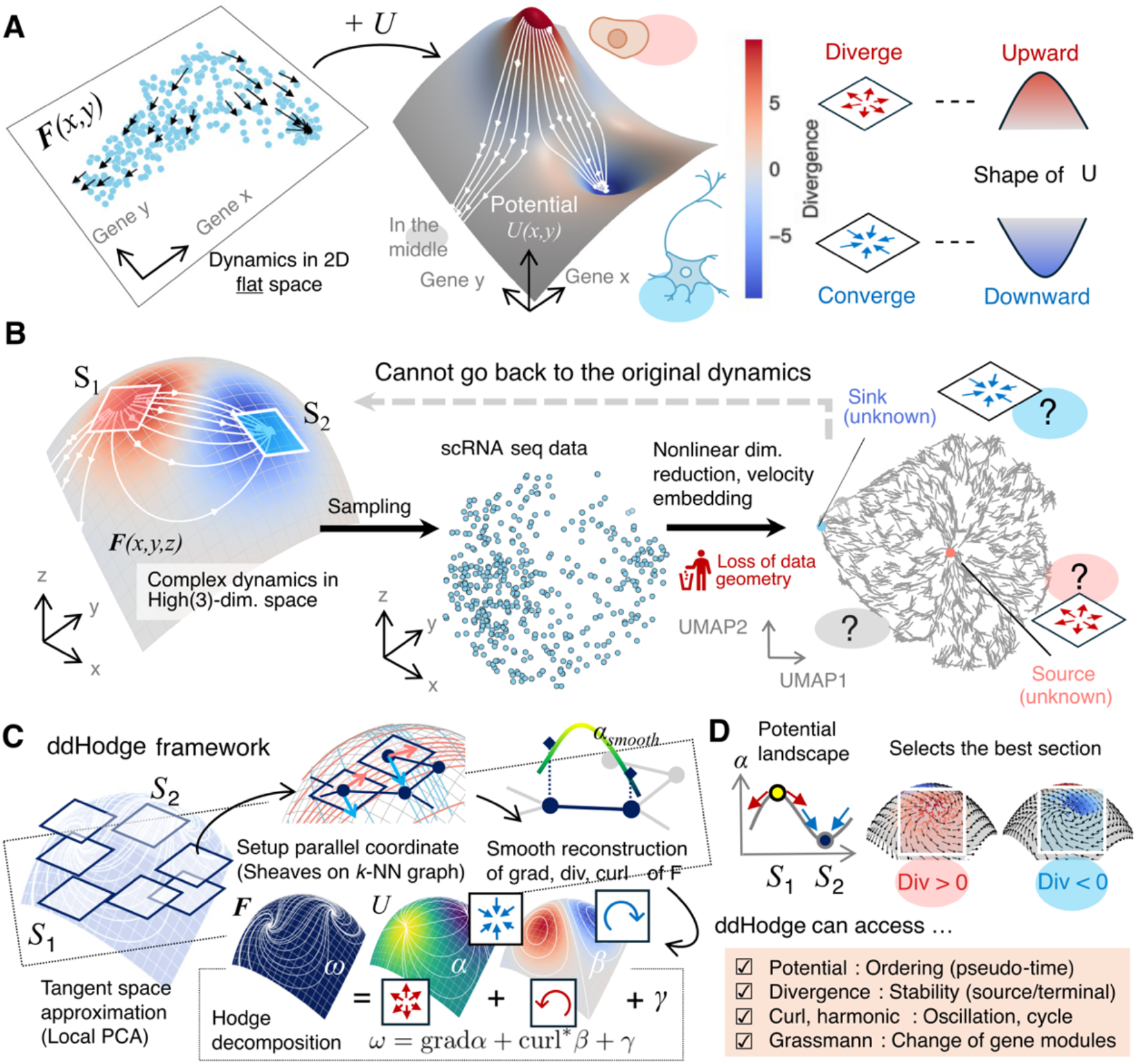
The ddHodge framework provides a method for analyzing unknown high-dimensional dynamics. **(A)** Shape of the landscape: The local shape of the landscape represents the local dynamics of cell states. The underlying dynamics of the cell state generates single-cell data (left), and the dynamics are represented by a potential landscape (middle). The shape of the landscape indicates the type of change in the cell state, such as diversification or convergence (right). A negative divergence (concave shape) indicates a stable state change in which velocities converge toward the center. A positive divergence (convex shape) indicates an unstable state change in which the cell state trajectories become more diverse. A divergence of zero (flat shape) indicates a change that occurs at a constant speed (i.e., no acceleration or deceleration). **(B)** Dimensionality reduction distorts dynamics. Single-cell data are sampled from unknown high-dimensional gene expression state space (left to middle). The sampled data are visualized in 2D by removing the geometric features of the high-dimensional data, such as the length and volume in the state space (right). This loss results in the loss of the shape of the landscape. The velocity embedding in the distorted space also inaccurately represents the dynamics of cell states. The closure of these boundaries in the visualized map did not occur in the original space. **(C)** ddHodge strategy: Reconstruct dynamics in the low-dimensional tangent space patches and combine these patches smoothly. The low-dimensional tangent spaces at each data point are estimated using a local PCA approach. A complete description of the vector field (components of Hodge decomposition: grad, div, and curl) is reconstructed smoothly along the near-parallel coordinate system designed in the curved space of the “data manifold” (the details are provided in the Methods). **(D)** ddHodge provides rich information about high-dimensional dynamics: the stability of states, the best section to visualize, and the changes of involved genes.

Reconstructing cell state dynamics after nonlinear dimensionality reduction involving data deformation is a considerably difficult problem^15–17^. Nevertheless, nonlinear dimensionality reduction is useful for visualization. We set the original high-dimensional (3D) dynamical system to have a region of instability (S1) with a large positive divergence and a region of stability (S2) with a large negative divergence (Figure 1B, left). However, in commonly used nonlinear dimensionality reduction methods such as UMAP^18^, the density of the data points in original space is equalized in the reduced space; as a result, geometric features such as length and volume are lost. This deformation affects the landscape shape, making it difficult to identify regions S1 and S2 existed in the original space (Figure 1B, right). Additionally, the velocity embedding approach^10,15^ used for visualizing velocity information can misrepresent dynamics by embedding high-dimensional velocity data with neighboring data points in the reduced space, potentially presenting nonexistent closed trajectories at the data boundaries.

To address the challenge of reconstructing high-dimensional dynamics, we propose a combination of two approaches. First, we introduce a high-precision vector field reconstruction method based on Hodge decomposition. A vector field on a manifold can be decomposed into three independent components: a gradient with divergence, a divergence-free rotational component, and a harmonic component as the residual (Hodge decomposition). The goal is to completely reconstruct these three components. Next, to make this method applicable in high-dimensional spaces, we consider approximating the Hodge components on a low-dimensional manifold (Figure 1C; dimensionality reduction preserving data geometry). By using local principal component analysis (PCA) to approximate tangent spaces^19^ and patchworking these spaces, we build a low-dimensional curved space embedded in the high-dimensional space (hereafter referred to as the data manifold) (Figure 1C, left). During the reconstruction process, we smooth the estimated values along the curved axes on the data manifold to compensate for noise and sample size limitations (Figure 1C, top), enabling the reconstruction of the potential, divergence, and curl in high dimensions (Figure 1C, bottom).

We implemented this framework as ddHodge.

ddHodge allows us to evaluate the temporal order and stability of data points on the basis of the potential and divergence, respectively. The periodic and oscillatory behavior of dynamic processes, such as the cell cycle and circadian rhythms, can be evaluated using the curl and harmonic components, independent of the gradient component. Even after dimensionality reduction, the Grassmann distance (the degree of bending when tangent spaces are patchworked) allows us to identify the timing of significant changes of dimensions (genes). The Jacobian matrix reconstructed on the data manifold can be used to select cross-sections of the gene expression space in which the trajectories significantly change. By extracting and visualizing these rich dynamic indicators (Figure 1D), we can identify genes associated with the diversification and convergence of dynamic cell states.

### ddHodge accurately reconstructs the true dynamics from partial velocity sampling

#### Reconstruction of the potential determines the height of the landscape

We evaluated the performance of ddHodge with synthetic data (Figure 2). The evaluation metric was the mean squared error (MSE) between the theoretical values of the predefined potential U(x, y) and the reconstructed values. Additionally, we assessed the ability of ddHodge to reconstruct the shape of the potential landscape (divergence) and compared it with that of conventional methods.

**Figure 2.**
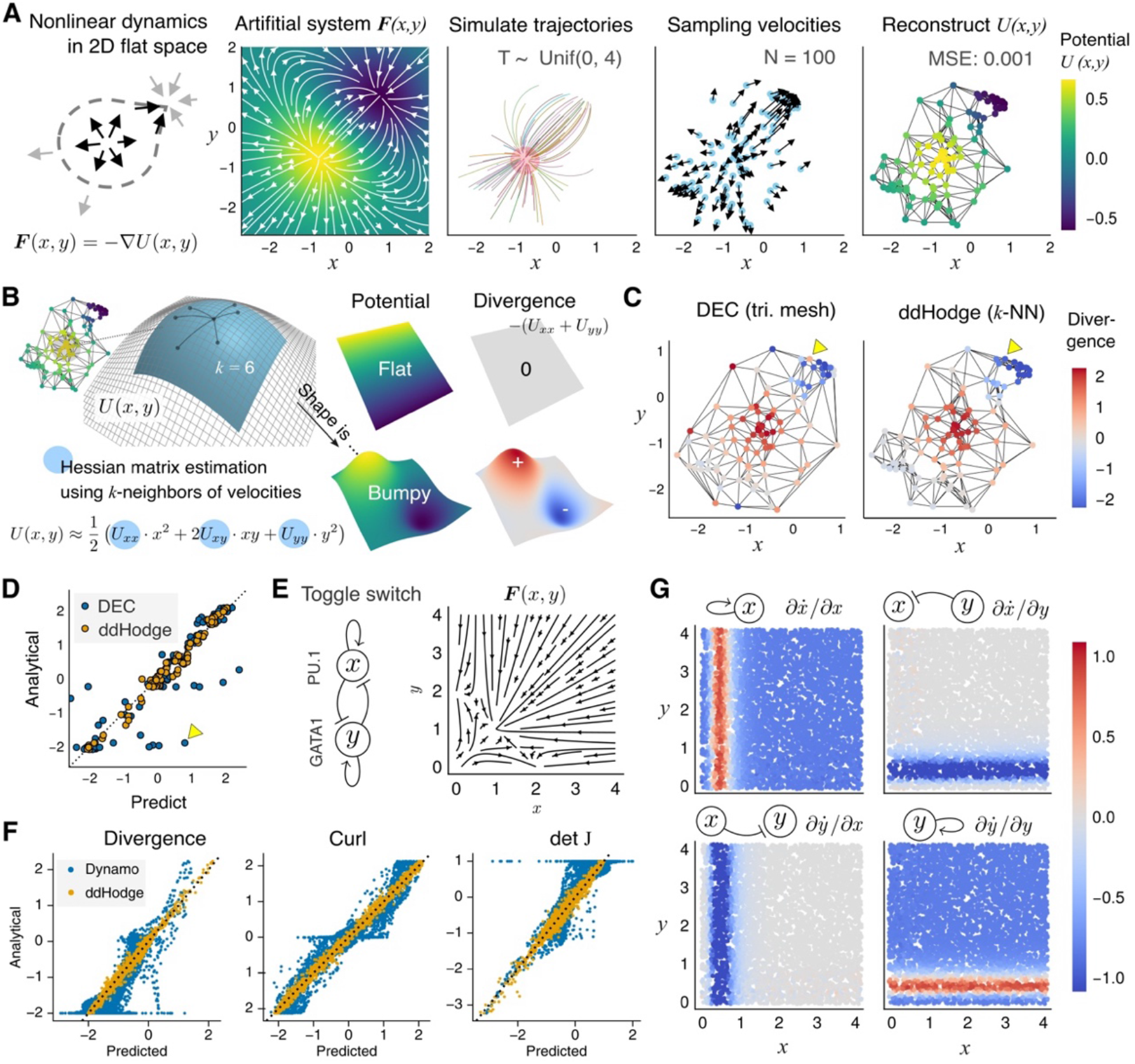
ddHodge accurately reconstructs dynamics from a partial sample of velocities. **(A)** Workflow of the simulation validation approach. We used the nonlinear system ***F****(x, y)*, which is completely determined by the gradient of a predefined potential *U(x, y)* in a low-dimensional flat plane (2D Euclidean space) (left). The initial values were set around the highest potential peak (2^nd^ to 3^rd^ panels). Using 100 data points sampled after random time intervals from 0 to 4 seconds (3^rd^ to 4^th^ panels), the potential values were reconstructed (right). **(B)** Recovering the shape of the landscape: The local shape of the landscape (Hessian matrix of U) can be recovered from the 2nd derivative of the piecewise quadratic functions fitted along the paths of the points to the *k* nearest neighbors. These two landscapes (middle column, top and bottom rows) have different shapes with the same potential values. The flat shape with zero divergence (top-right) indicates constant changes of states, whereas the bumpy shape with large positive and negative divergence (bottom-right) indicates unstable states. **(C)** Comparison with the DEC method in terms of the divergence reconstruction performance: DEC with Delaunay triangular mesh vs. ddHodge with a *k*-NN graph. The yellow arrowheads at the sampling boundaries indicate the marked difference in the divergence estimated by the DEC and ddHodge methods. **(D)** Direct comparison of the performance of the two methods: The x-axis is the predicted value of the divergence by the DEC and ddHodge methods. The y-axis is the analytical (true) divergence of ***F***(x, y). The yellow marker indicates the estimated values at the arrowhead on the left side of Figure 2C. **(E)** Toggle-switch model: A nonlinear mathematical model of cell differentiation for generating multistable states is visualized with streamline plots. **(F)** ddHodge produces substantially more accurate reconstruction results than does Dynamo: the divergence, curl, and Jacobian (detJ) results obtained using Dynamo and ddHodge are shown. **(G)** Accurate recovery of the Jacobian matrix elements with ddHodge. Each panel indicates the estimated value of the Jacobian matrix elements (J_11_ J_12_ and J_21_ J_22_).

To evaluate the performance of ddHodge in reconstructing dynamic information without dimensionality reduction, we conducted simulations of a dynamical system in which the trajectory of each point on a 2D plane was determined solely by the gradient of the potential U, defined as ***F***(x, y) = -∇U(x, y) (Figure 2A, 1st and 2nd panels, Methods). We randomly initialized points near the peak of the potential in the lower left region and simulated their trajectories for random durations between 0-5 seconds, obtaining 100 pairs of position and velocity data (Figure 2A, 3rd and 4th panels). When ddHodge was applied to estimate the potential via the least squares method introduced in HodgeRank^13^, the reconstructed values agreed well with the theoretical values (Figure 2A, 5th panel; MSE: 0.001). This result demonstrates that the proposed method can provide high-precision potential reconstruction results even with small datasets and partial sampling of data points.

#### Reconstruction of the shape of the landscape indicating stability

We next leveraged the geometric interpretation of the estimated potential to reconstruct the shape of the potential landscape. We viewed the least squares potential estimation method as fitting a piecewise quadratic function to the edges of a graph with given slopes at the ends (Supp. Notes). This approach can be extended to estimate the shape of the potential function (the Hessian matrix H of U). The shape is approximated by collecting fitted piecewise quadratic functions of each cross-section to the neighboring vertex (Figure 2B, Methods). The shape of the potential indicates whether the cell state traverses the landscape at a constant speed (flat slope; upper case in Figure 2B) or experiences acceleration, deceleration, or passes through branching points (changing slope; lower case in Figure 2B).

Next, we verified the reliability of ddHodge’s results. We compared the divergence reconstruction performance of ddHodge with that of the discrete exterior calculus (DEC) method^20,21^, which is equivalent to the finite element method (FEM), a well-established numerical analysis method (Figure 2C). The results showed that ddHodge achieved a reconstruction accuracy comparable to or better than that of the DEC method, even when the data boundary, where the DEC method had lower reproducibility, was excluded (yellow triangles in Figures 2C-D) (Figure 2D, DEC: MSE = 0.502, ddHodge: MSE = 0.026).

#### ddHodge outperforms Dynamo in terms of reconstruction performance

Next, to evaluate the reconstruction performance of ddHodge, we compared ddHodge with Dynamo, which has similar vector field reconstruction capabilities. In this comparison, we used simulation data from the toggle-switch model, a 2D nonlinear differential equation model (Figure 2E). We compared the reconstruction results of the two methods using *N =* 5,000 uniformly sampled pairs of position (x, y) and velocity ***F***(x, y) data from the toggle-switch model^12^. The curl was reconstructed using the ddHodge’s divergence of the *dual* velocity fields (Methods, Supp. Notes). The results showed that ddHodge significantly outperformed Dynamo in all metrics: the divergence (ddHodge vs. Dynamo: 0.0027 vs. 0.1142), curl (0.0027 vs. 0.0742), and Jacobian (0.0051 vs. 0.1050) (Figure 2F). These results may be attributed to the differences in the design principles of these two methods, i.e., ddHodge has the same design principles as the exact calculation methodologies used in conventional numerical analysis techniques, such as DEC, whereas a machine learning approximation technique is employed with Dynamo^22^.

Furthermore, the Jacobian matrix J can be reconstructed from the accurately reconstructed Hessian matrix and the curl (Methods, Supp. Notes). We then evaluated the applicability of the proposed method for interaction analyses of gene pairs using J. The clear red and blue bands in the reconstruction results of each element of the Jacobian matrix (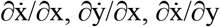, and 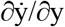 where 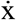 represents dx/dt) faithfully reproduced the regions that respond sensitively to perturbations in genes x and y (Figure 2G).

These results demonstrate the ability of ddHodge to reconstruct unknown nonlinear dynamics with high accuracy and efficiency, even from partially sampled data.

### Approximating low-dimensional dynamics on the data manifold

Next, we aimed to develop a dimensionality reduction method suitable for high-dimensional dynamical systems (Figure 1C, top). For example, the surface of a sphere in three dimensions can be represented as a collection of small two-dimensional planes (Figure 3A, top). Based on this concept, we hypothesized that by reconstructing and stitching dynamics within local lower-dimensional spaces (tangent spaces) using a local PCA approach^19^, both dimensionality reduction and faithful reproductions of high-dimensional dynamics could be achieved while preserving the geometry of the data manifold. The number of parameters in the Hessian matrix estimation process is reduced to *m*^2^ << *d*^2^, where *d* is the number of genes and *m* is the dimension of the tangent spaces. We also considered smoothing (regularization) along the curved axes on the data manifold. In the smoothing process, the axes of adjacent tangent spaces should be matched (Figure 3A, middle; *x*_*u*_ and *x*_*v*_, *y*_*u*_ and *y*_*v*_); therefore, using idea of parallel transport in differential geometry^19,23,24^, we developed a method for designing near-parallel coordinate axes for high-dimensional data using a discrete connection Laplacian (sheaf Laplacian) ^24–27^ (Figure 3A, bottom; Supp. Notes).

**Figure 3.**
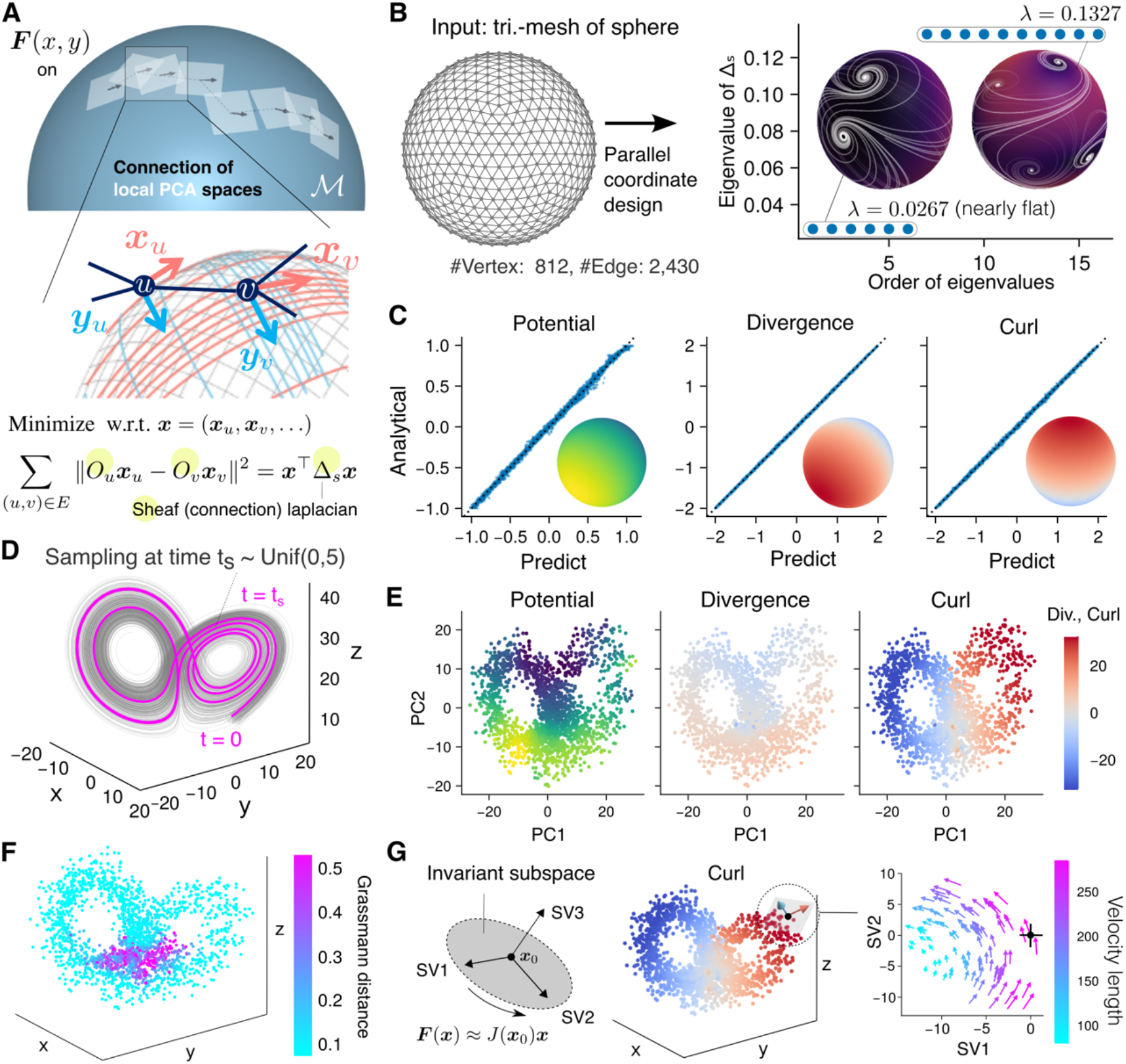
Rebuilding high-dimensional dynamics by patchworking lower-dimensional local dynamics. (**A**) Designing near-parallel coordinates on curved spaces: The surface of a sphere, as an example of a curved 2D space embedded in 3D, is represented by small patches of flat tangent planes (top). The arrows (local x- and y-axes, x_u_ and y_u_) to the adjacent tangent planes involves the problem of parallel transport (middle). The solution is to find the minimum eigenvalue of a sheaf Laplacian (bottom). (**B**) Near-parallel coordinates on a sphere: A triangulated spherical mesh is used as the input graph. The eigenvalues of the sheaf Laplacian reveal the degrees of freedom in designing near-parallel axes. The insets show two examples of designed near-parallel coordinates. (**C**) Near-perfect reconstruction of dynamics on a sphere: Scatter plot of *N* = 2,000 random points comparing the reconstructed (horizontal) and analytical (vertical) values of the potential, divergence, and curl on a sphere. The insets show the analytical values, with the peaks of the potential and divergence on the west and east sides and the peaks of the curl on the north and south sides. (**D**) Sampling scheme from the Lorenz system: *N* = 2,000 points and velocities sampled at random intervals from 0 to 5 seconds, starting from a random initial point. (**E**) Lower-dimensional representation of Lorenz system: The dynamic indices (potential, curl, and divergence) reconstructed in estimated tangent planes via a local PCA approach, visualized on a global (i.e., including all sampled points) PCA plane. (**F**) Grassmann distance reveals hinge points: The distance indicates the folding angle when patching local PCA planes. (**G**) Schur decomposition: The section that highlights critical changes at a specified point is selected. The local dynamics can be approximated by the Jacobian matrix J. The rotating dynamics around x_0_ (point with the highest curl) on the right side are highlighted by projecting the velocities onto the section spanned by Schur vectors 1 and 2.

First, we present a simple example of designing near-parallel coordinates on a sphere (Figure 3B). We used the graph structure of a triangulated mesh as the input. By adopting the eigenvector corresponding to the smallest eigenvalue of the connection Laplacian of the input graph, we obtained a near-parallel axis that is consistent^23,24^ on the surface (Figure 3B right). Additionally, from the distribution of the obtained eigenvalues, we found that the best near-parallel coordinates that minimize the deviation from perfect parallel transport on this spherical mesh can be designed with 6 degrees of freedom. Furthermore, there was a gap^28^ in the spectrum of the eigenvalues from the 7th value onward, corresponding to the doubling of the number of visible vortices, known as singularities^23^.

Next, using velocities sampled from the artificial dynamics simulated on a sphere (Methods), we reconstructed the dynamics with smoothing on the data manifold (Figure 3C). The potential and divergence were reconstructed with peaks at the east and west ends of the sphere, and the curl was concentrated at the north and south ends, reproducing values approximately identical to the theoretical values (Figure 3C; inset images, MSEs: 1.0×10^−3^, 4.0×10^−4^, and 1.0×10^−3^, respectively). These results indicate that both the unknown low-dimensional manifold and the dynamics on the manifold can be accurately recovered from the sampled data.

### ddHodge enables qualitative interpretation of unknown complex systems

Next, we validated the interpretability of the lower-dimensional approximation of high-dimensional dynamics using the Lorenz system (Figure 3D). The dynamics of the Lorenz system are characterized by trajectories that irregularly move between two disks in three-dimensional space. Therefore, we attempted to approximate the dynamics in the space of two joint planes as a data manifold and capture the characteristics of the original dynamics (Figure 3E). As expected, the main characteristics of rotational behavior were identified by the positive (anticlockwise) and negative (clockwise) curl values on the left and right sides of this space. Additionally, the acceleration in the trajectory in the lower half of the space and deceleration in the upper half of the space were confirmed by the sign of the divergence value. In summary, as indicated by the smaller divergence values compared with the curl values, the dynamics of this system were considered dominated by rotations rather than gradients. Furthermore, the Grassmann distance was used to identify the planar region of the trajectory (cyan) and the hinge region at the center (magenta; points of dimensional change) (Figure 3F).

The representative variables (genes) driving the dynamics can be identified by the Schur decomposition^15,29^ of the Jacobian matrix (Figure 3G, left). Here, we used the ddHodge results in three dimensions without approximating the Lorenz system in 2D space (Figure 3G, middle). As expected, prominent rotational dynamics were observed on the SV1-SV2 plane at the point at which the curl has its maximum value (Figure 3G right). These results indicate that the Schur decomposition of the Jacobian matrix is effective for identifying the cross-section of the space containing the representative dynamics nearby specified point (in this example, the cross-section along the disk).

These results demonstrate the utility of ddHodge in inferring the behavior associated with the original dynamics, even in the approximate reconstruction of the high-dimensional dynamics.

### The data analysis framework of ddHodge provides a global view of embryogenesis

Next, the scRNA-seq data were analyzed using ddHodge to quantify the potential, which represents the order of cell state transitions, and divergence, which reflects the stability of each cell state (Figure 4A). In this analysis, we focused on VASA-seq data, which provide highly accurate velocity information due to their high degree of intron coverage ^6^. By leveraging this dataset, we attempted to elucidate the hierarchical differentiation sequence of cell populations and the changes in the stability of the cell state for each cell lineage during the embryogenesis.

**Figure 4.**
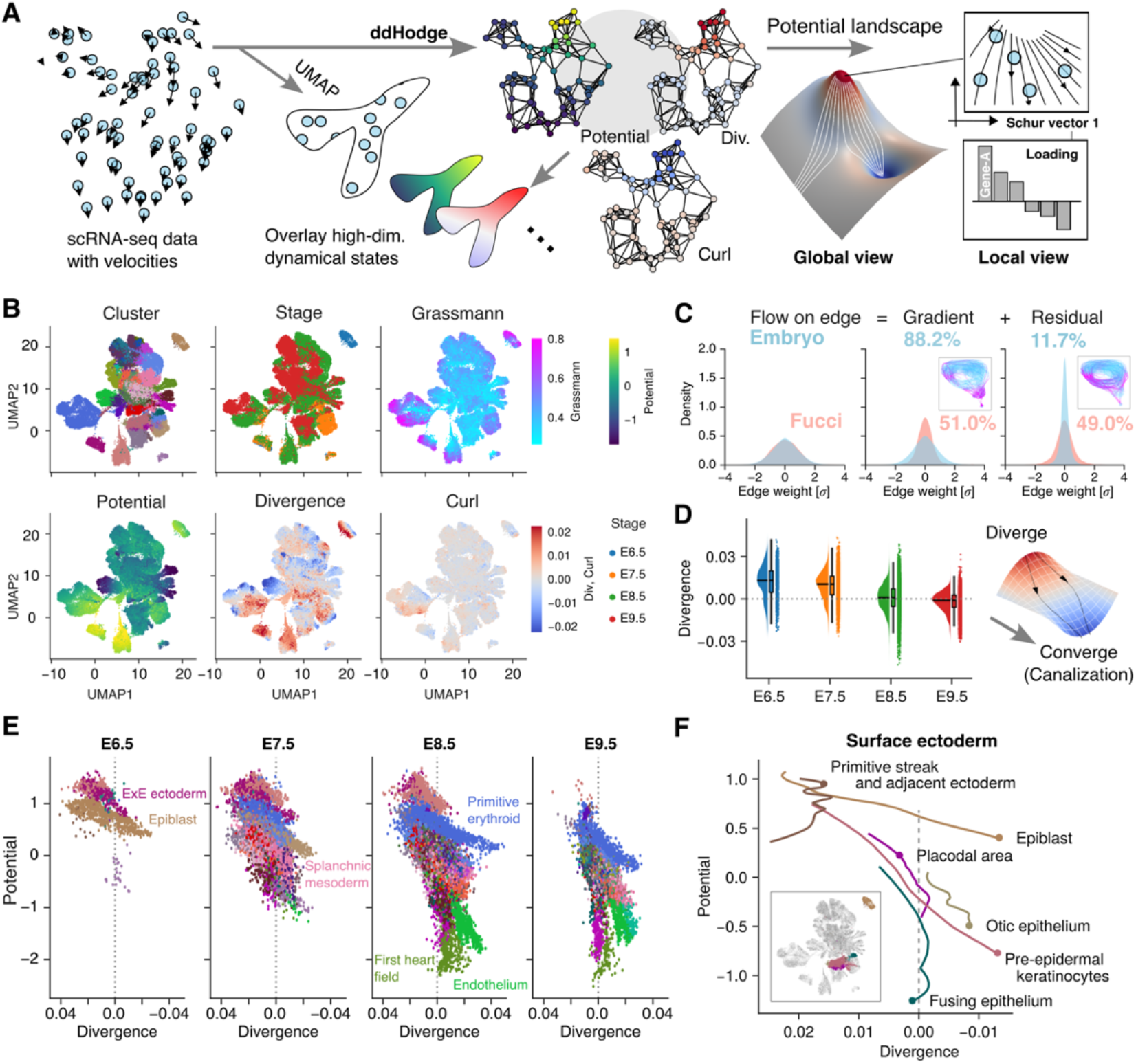
Potential landscape provides a succinct global view of gene expression dynamics in embryogenesis (A) scRNA-seq data analysis framework: ddHodge takes scRNA-seq data paired with velocity information (e.g., RNA velocity) as input (left). The recovered dynamic indices are overlaid on visualizations such as those obtained via UMAP (middle), providing a global view of high-dimensional dynamics through landscape representations. From this global view, cells of interest can be determined by the extracted dynamical indicators (right), allowing detailed investigation of specified regions (local view) in which gene expression state changes are prominent, as well as the genes involved. **(B)** Dynamical indicators extracted during mouse embryogenesis. In addition to clusters and embryonic stages, four dynamical indicators extracted by ddHodge are shown. VASA-seq data from mouse embryos from E6.5 to E9.5 were analyzed here. **(C)** The majority of the gene expression dynamic indicators can be explained by a potential landscape: The value of the gradient and the residual of the divergence-free (curl + harmonic) components of recovered dynamics data are shown. For comparison, the results of ddHodge based on the FUCCI dataset, which is expected to have a dominant cyclic component, are also shown. **(D)** Changes in the stability of the cell state during development: The boxplot (Q1, Q3 and median) of global change in the divergence from E6.5 to E9.5 is shown. The right image shows a canalization-type landscape, where cells transition from diversification (unstable) to convergence (stable). **(E)** Summarized dynamics of embryonic development using the height and shape of the potential: For each embryonic day, the vertical axis shows the potential, which indicates the temporal order of the state change, and the horizontal axis shows the divergence (axis is flipped), which represents the stability of the gene expression state. **(F)** Simplified landscape view of development: A potential-divergence plot of the surface ectoderm lineages extracted from (E), represented as thin lines. The inset shows the positions of the extracted clusters in the UMAP representation.

The VASA-seq data contained *N* = 46,124 cells from mouse embryos in the stage of E6.5–9.5. Using the gene expression levels and RNA velocity as inputs, the components of the Hodge decomposition—potential, divergence, curl, and Grassmann distance—were reconstructed and mapped to a low-dimensional UMAP representation using different colors (Figure 4B). We found that the Grassmann distance remained low across the dataset (median value: 0.43), while the reduced dimension was *m* = 4. This result suggests that the trajectories of the gene expression dynamics associated with embryonic development reside in a four-dimensional manifold. We also found a correlation between the potential value and developmental stage, i.e., the potential progressively decreases as development progresses, indicating its utility as a pseudotime indicator^12^ (Supp. Fig. S1). Similarly, the divergence became increasingly negative as development progressed. Notably, the value of the curl, which reflects cyclic/oscillatory dynamics such as those associated with the cell cycle, was substantially lower than that of the divergence. These findings suggest that the majority of the gene expression dynamics during embryogenesis can be described by a gradient system governed by a potential landscape.

To further confirm that the dynamics of embryogenesis is mainly driven by a gradient system, we quantified the proportion of cyclic or periodic components separated from the gradient component (Figure 4C). In this dataset, the gradient component accounted for 88.2% of the gene expression dynamics, whereas the divergence-free cyclic component accounted for only 11.7% of the gene expression dynamics. In contrast, in the control dataset of FUCCI-labeled cells^30^, which specifically represents the progression of the cell cycle, the cyclic component accounted for 49.0% of the dynamics (Figure 4C, magenta; Methods). These findings suggest that embryonic gene expression dynamics can be explained mainly by the potential rather than by periodic components such as the cell cycle. This underscores the importance of understanding the topography of the landscape to elucidate the mechanistic underpinnings of embryogenesis.

To further elucidate the biological significance of this divergence, we analyzed its changes over the course of embryogenesis (Figure 4D). During the early stages (E6.5–E7.5), most cells exhibited positive divergence, indicating an undifferentiated and unstable (divergent) state. In contrast, from E8.5 to E9.5, the divergence values became increasingly negative, suggesting that the gene expression states of the cells tended to converge and stabilize as development progressed. This finding indicates that early embryonic cells exhibit a high degree of plasticity, which subsequently decreases as differentiation becomes more defined and cell states stabilize.

Thus, we attempted to describe the dynamics of embryogenesis using only the potential and divergence (Figure 4E). The epiblast, which possesses pluripotency with high divergence, maintained its cellular identity with a shallow slope (gradient) between E6.5 and E7.5 but was no longer detectable after E8.5. In contrast, cell populations that had diverged into distinct lineages, including the splanchnic mesoderm, emerged beneath the epiblast from E7.5 onward and progressively descended along the landscape toward E9.5, differentiating into more stable cardiac and epithelial lineages. These findings suggest that we successfully visualized the fate decision process that occurs during development according to the height and shape of the potential landscape during gastrulation.

Next, we focused on the surface ectoderm lineage (Figure 4F). The surface ectoderm, a nonneural ectoderm, gives rise to the epithelium and epidermis of sensory organs. Even at E8.5, pre-epidermal keratinocytes appeared at a higher positions in the plot (potential > 0.5) than the placodal area, suggesting that the epidermal lineage may be specified earlier than the epithelial lineage^31^. Furthermore, pre-epidermal keratinocytes and the otic epithelium exhibited stable cell states at E9.5, as indicated by the highly negative divergence values (approximately -0.01). In contrast, the divergence of the fusing epithelium, which exhibits gene expression characteristics associated with the amniotic ectoderm, remained close to zero, indicating that the cell state changed with a constant speed, which is a unique property of this type of cell.

Taken together, our results demonstrate that the developmental trajectory of the surface ectoderm lineage is consistent with a recent single-cell study of embryogenesis^32,33^. Moreover, we successfully quantified the potency of differentiation at each developmental stage using the potential and divergence.

### ddHodge enables the extraction of critical timing information and genes related to cell state changes

As nonlinear dimensionality reduction is not performed in ddHodge, the genes that drive cell state changes can be tracked. We thus focused on individual points of interest (POIs) at which cell states undergo significant changes and investigated the local dynamics (Figure 5).

**Figure 5.**
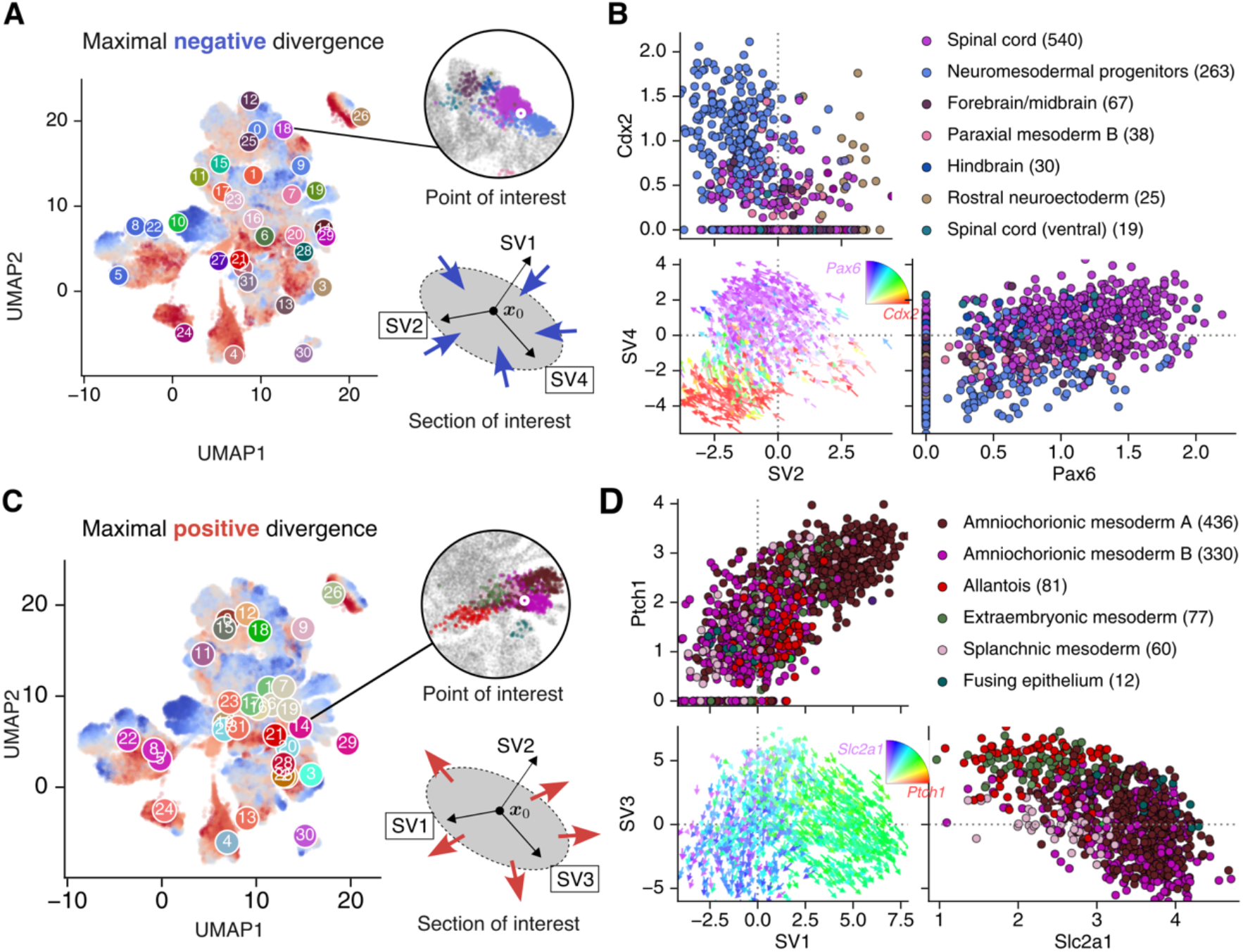
Cross-sections of the gene expression state space to elucidate the local dynamics of embryogenesis (A) (Case 1) Stabilization of a cell state with a highly negative divergence in axon elongation: 2,000 cells surrounding the selected spinal cord cells that had minimal negative divergence values in cluster 18, including the NMP and forebrain/midbrain cell types, were selected (inset circle; enlarged image of the UMAP plot near the POI). From the Schur decomposition of the Jacobian matrix at the POI, the two dimensions (plane) with the minimal negative eigenvalues were determined. **(B)** The velocity of the 2,000 selected cells projected onto the identified SV2-SV4 cross-section, along with the genes exhibiting significant changes within this plane, are shown. The cell type annotation is shown for the top six cell types, and the numbers in parentheses indicate the number of cells in this plot. **(C) (Case 2)** Diversification of the cell state with a highly positive divergence in the amniochorionic mesoderm: 2,000 cells surrounding amniochorionic mesoderm cells that had maximal positive divergence values in cluster 14, including allantois, extraembryonic mesoderm, and splanchnic mesoderm cell types, were selected. The two Schur vectors with the maximal positive eigenvalues were determined. **(D)** The velocity of the 2,000 selected cells projected onto the identified SV1-SV3 cross-section, along with the genes exhibiting significant changes within this plane, are shown. The expression ratios of the two selected genes in (C) and (D) are shown in rainbow colors (inset, quarter disc).

To identify cells that become more stable, namely, those with negative divergence values, we selected the POI with the smallest divergence within cluster 18 (Figure 5A, top right). The Schur decomposition of the Jacobian matrix obtained for this POI revealed a remarkable state change in the SV2-SV4 plane, corresponding to the largest negative eigenvalues. To determine the most related genes in this section, we identified candidate genes using the correlation coefficients between the divergence and gene expression levels for 1,000 neighboring cells in the POI (Supp. Fig. S2). The velocities projected onto the SV2-SV4 planes demonstrated that the cell states converged and stabilized (negative divergence); furthermore, *Pax6* expression was identified^34^ (Figure 5B, Supp. Fig. S2). Additionally, *Nrcam* and *Robo2*, which are genes related to axon guidance^35^, exhibited strong negative correlations with the divergence values, similar to *Pax6*, suggesting their involvement in the stabilization of cell states during axial elongation. In contrast, another cell population annotated as neuromesodermal progenitors (NMPs) expressing *Cdx2* was positively correlated with divergence (Figure 5B, top and bottom left panels, Supp. Fig. S2). *Cdx2*, which is regulated by multiple extracellular signals, has been reported to play a role in maintaining bipotent NMP populations^36,37^. These findings suggest that the mechanism maintaining undifferentiated NMPs may involve destabilization of cell states through *Cdx2* expression. In summary, through this analysis, we identified several genes associated with the spinal cord lineage involved in regulating cell state stability.

To identify genes associated with cell state diversification, we selected another POI with the highest divergence from cluster 14 (Figure 5C, top right). The neighboring cells of this POI primarily consisted of two amnion-like cell groups (annotated as A and B) (Figures 5C, D). The Schur decomposition of the Jacobian matrix at the POI revealed two branches along the SV1 axis, with one group diverging to the right (A) and the other to the left (B). Since the loading vector of the Schur decomposition can be used to identify genes that are strongly associated with the selected axis (Supp. Fig. S3), we attempted to identify the genes with the highest weights in the loading vector. *Ptch1* presented the highest loading value along the SV1 axis, suggesting the important role of this gene in A/B fate determination (Figure 5D, top). *Ptch1*, the receptor of Sonic hedgehog (Shh), has been reported to be expressed in the allantois, leading to the arrest of the epithelial-mesenchymal transition and the progression of morphogenesis^38^. Therefore, only group A, which expresses *Ptch1*, may undergo de-repression via Shh signaling and transition to a distinct cell state. Additionally, *Slc2a1*, which encodes a glucose transporter of Glut1, presented the highest loading value on the SV3 axis; however, its expression level increased with developmental progression in both A and B. In summary, this axis-based approach can provide a biological interpretation similar to that of PCA.

## Discussion

Here, we developed ddHodge as a method for reconstructing the original vector field using pairs of sampling points and their velocities obtained from an unknown system. By utilizing RNA velocities as inputs, ddHodge extends the capabilities of existing methods by allowing evaluations of both the direction of cell state changes and the stability of each state, which indicates a potency of change, which has not previously been evaluated. We further confirmed that the majority of the gene expression dynamics in embryogenesis can be described as a gradient system (i.e., a potential landscape), which was previously unclear. Furthermore, we quantified the difficulty of lineage transitions from the divergence values, complementing traditional trajectory analyses of cell fate conceptualized by the epigenetic landscape. Additionally, we identified genes that influence the stability of the cell state.

In this study, we demonstrated that the potential and divergence reconstructed from RNA velocity data can serve as indicators for identifying points at which cell states converge and diversify. Furthermore, ddHodge can be used to extract cross-sections of high-dimensional space and identify key regulatory genes that govern local dynamics. Since this framework is general and requires only a weak assumption that the velocity is sampled from an autonomous system, ddHodge can be broadly applied to various types of velocity data, including chromatin velocity data, protein velocity data, time-resolved measurements of cellular positions, and even simulation data from theoretical models of high-dimensional dynamical systems. Notably, as evidenced by the constructed interpretations of the high-dimensional dynamic indicators in both the simulations with the Lorenz system and the application to the RNA velocity data, the approximated velocity field on the data manifold does not necessarily need to precisely capture the true underlying dynamics.

ddHodge reconstructs the fundamental components of vector fields, including the potential, divergence, curl, and harmonic by overcoming several difficulties associated with analyses performed in high-dimensional spaces. For example, in the DEC method, these difficulties include constructing the mesh, evaluating areas and volumes, which are required to determine the densities and constructing discrete Hodge stars^20,21^. With our approach, we could accurately estimate the divergence by applying previously intractable numerical analysis methods to higher-dimensional spaces without considering boundary conditions. Additionally, we developed a method for estimating the curl in spaces of three or more dimensions. To address concerns regarding the reliability of the results due to noise and limited sample sizes—challenges inherent in data analysis—we incorporated a dedicated smoothing strategy in ddHodge. However, due to computational constraints, we have not yet produced a strict reconstruction of the curl component with ddHodge; instead, an approximation strategy that recovers the rotational features only within the PC1-2 plane is employed in the current version of ddHodge. Further interpretation of divergence-free periodic components may be feasible by developing efficient methods for computing curl by Hodge duals in high-dimensional spaces and establishing appropriate interpretative frameworks, such as optimal cycles^39^, to obtain a succinct representation of the harmonic components in higher-dimensional spaces.

Owing to its scalability to high-dimensional data without the need for mesh construction, this method has potential as a tool for supporting data analysis involving high-dimensional complex dynamic systems. The remaining fundamental challenges are interpretating the recovered dynamic indicators in the context of existing knowledge and experimentally validating the proposed method.

## Methods

### ddHodge workflow

This section outlines the ddHodge workflow. Detailed explanations and illustrations are provided in the Supplemental Notes. We assume that the input data consist of *d*-dimensional points *χ* = {**x**_1_, **x**_2_, …, **x**_N_} and their corresponding velocities 𝒱 = {**v**_1_, **v**_2_, …, **v**_N_}. Additionally, a *k* - nearest neighbors (k-NN) graph *𝒢* = (*V, E*) is constructed from *χ*.

### Approximation of a low-dimensional data manifold

First, PCA is performed using the *k* -nearest neighbors of each point **x**_*i*_ to obtain the orthonormal basis 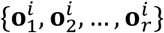 of the tangent vector space *T*_*i*_*ℳ*, where *r* ≤ *k*. Since the tangent spaces of neighboring vertices generally have different subspaces, we construct a discrete connection Laplacian *Δ*_*s*_ using sheaves on graphs to align the local coordinates within the tangent spaces. The alignment is performed by minimizing the following loss function *L* over the set of tangent vectors **z**_*i*_ ∈ *T*_*i*_*ℳ*:

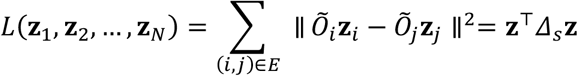

where **z** is the vector formed by stacking **z**_1_, **z**_2_, …, **z**_N_ vertically. The matrices Õ_*i*_ and Õ_*j*_ are obtained from the singular value decomposition 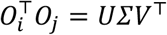, where 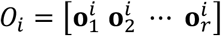 and Õ_*i*_ = *U*^⊺^, Õ_*j*_ = *V*^⊺^.

The eigenvector associated with the minimum eigenvalue of *Δ*_*s*_ is the minimizer of *L*.

### Potential estimation

The potential **u** ∈ ℝ^*N*^ corresponding to the gradient component of the vector field is estimated using the gradient operator δ_0_ = grad, which is an *M* × *N* matrix, and the edge weights **w** ∈ ℝ^*M*^ of graph *𝒢*. The estimation is performed via the least squares method as follows:

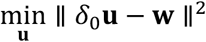

where *N* and *M* represent the number of vertices and edges in *𝒢*, respectively. In this study, the edge weights *w*_(*i,j*)_ are defined as:

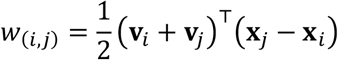

In this definition, the integral of the flux is approximated along the edge direction and is equivalent to fitting a quadratic function to the cross-section of potential function *U* (see Supp. Notes for more details).

### Hessian matrix estimation

For a point **x**_*i*_, let **η** = **x**_*j*_ − **x**_*i*_ be the vector indicating the direction from this point to a neighboring point **x**_*j*_. The second-order directional differential coefficient of the fitted quadratic function is given by:

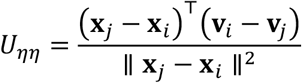

The Hessian matrix *H* in the original coordinate system can be obtained via a quadratic form. For example, in two dimensions of x-y plane, the relationship is expressed as:

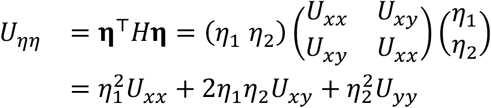

By gathering the known second-order differential coefficients such as *U*_*ηη*_ in independent directions *φ, ψ*, …, the linear equation can be constructed to estimate the elements of *H*.

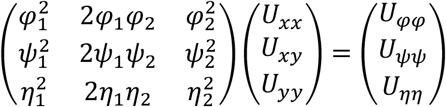

We denote the above equation as *C***β** = **y**. The divergence is obtained from −tr*H* after estimating *H*. In practice, the problem is better solved as an optimization problem min_**β**_ ∥ *C***β** − **y** ∥^2^ to mitigate issues in data analysis, such as those related to noise and the limited sample size. Additionally, since the number of Hessian matrix elements (number of estimated parameters) increases on the order of *d*^2^, the Hessian matrix is estimated in the reduced-dimensional space.

### Smoothing over graph

To enable the regularization of the estimation process between neighboring vertices, we merged the individual Hessian matrix estimation problems as a single linear equation:

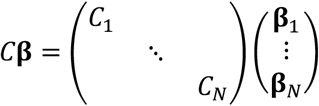

This allows us to introduce a penalty (Dirichlet energy on the graph) to the squares of the differences **β**_*i*_ − **β**_*j*_ between adjacent vertices as:

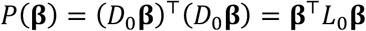

Here, assume that the *p* parameters are arranged in a certain order as 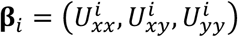, such that the differences between each parameter can be individually evaluated using the extended gradient matrix *D*_0_ = δ_0_ ⊗ *l*_*p*_ with the Kronecker product and 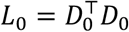.

The optimization problem with this regularization term is then solved as:

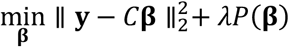

with some small constant λ > 0.

Given the large *Np* × *Np* size of the sparse matrix *L*_0_, efficient solvers such as the Krylov subspace method are recommended.

### Curl estimation

For differential forms in the Euclidean space ℝ^2^, the relationship between div and curl is given by:

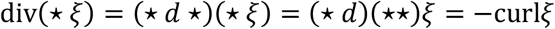

where *d* is the exterior derivative and ⋆ is the Hodge star operator. Therefore, the curl can be approximated by the divergence of the *dual vector field* ⋆ ξ, which is obtained by rotating the velocity by *π*/2 in the PC1-2 plane.

### Jacobian matrix estimation

According to the theory of Hodge decomposition, a vector field can be expressed as the sum of a gradient part −∇*U* and a divergence-free residual part *r*:

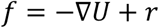

Here, let *D* be the operator that gives the Jacobian matrix of *f*. Since *D* is linear and *D*∇*U* gives the Hessian matrix *H* of *U, D*(*f*) has the following decomposition:

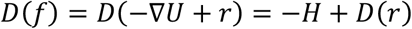

If *r* consists of only rotational components (i.e., there are no holes in the space), *D*(*r*) can be expressed using the matrix *Q*, representing a *π*/2 rotation in the PC1-2 plane, as 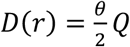. Here, *θ* is the magnitude of the rotation, which is calculated from the previously derived curl.

### Software implementation

ddHodge was implemented in Julia, a high-performance scientific programming language. The computation time depends on the number of analyzed cells and the dimensionality of the components after the reduction process is performed. The most computationally intensive parts are computing eigenvalues and solving least-squares problems with large sparse matrices. To optimize these calculations, we employ the Krylov subspace method (KrylovKit.jl), which efficiently handles sparse matrices by leveraging repeated matrix-vector multiplication operations. With GPU acceleration (we used Nvidia RTX A2000 12 GB), the computation takes approximately 35 seconds for 46,124 cells with the parameter: *k=12* of *k*-NN graph and *r=4* of reduced dimension, and approximately 2 minutes for another dataset with 98,316 cells with the same parameter (Figures 4-5).

### Dynamical system setup

#### Gradient system in the 2D plane

To create a gradient system in a 2D plane, we use the Gaussian function as follows:

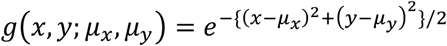

We constructed a potential function with peaks and valleys as follows:

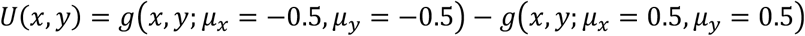

Using this potential function, we simulate the differential equation **F**(x, *y*) = −∇*U*(x, *y*). The simulation is performed by adding noise (Gaussian noise with a standard deviation of 0.2) to the initial positions at the center of the potential peak and running random simulations for 0 to 4 s using the Runge–Kutta method. We sampled *N* = 100 points and their velocities.

#### Toggle-switch model

The toggle-switch model is defined by the following set of differential equations:

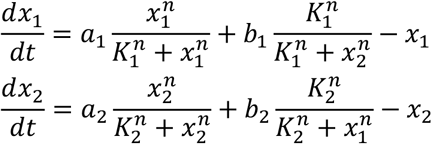

The parameters are set to *K*_1_ = *K*_2_ = 1/2, *n* = 4, and *a*_1_ = *a*_2_ = *b*_1_ = *b*_2_ = 1. The same simulation result from Dynamo’s paper was reproduced by following the instructions at the following link: https://github.com/aristoteleo/dynamo_submission.

#### Lorenz system

The Lorenz system is defined by the following set of differential equations:

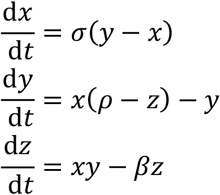

The parameters are set to *σ* = 10, *ρ* = 28, and *β* = 8/3. The simulation is performed by initializing (x, *y, z*) = (10,10,10) with added Gaussian noise *N*(3,0) and sampling points after a random time *T*, which was generated from a uniform distribution *T* ∼ *U*(0.5,5).

### scRNA-seq data analysis

For RNA velocity analysis, spliced and unspliced read counts were obtained from publicly available processed data in text format (VASA-seq) or loom files (FUCCI). Velocities were estimated using scvelo (version 0.3.2). For cell type annotation, the scRNA-seq data of mouse embryos ^32^ were used as training data for Celltypist (version 1.6.3), and the cell type labels were given for the VASA-seq data. The analysis using ddHodge was performed with a *k*-NN graph where *k* = 12 and the reduced dimensionality was *r* = 4. The detailed methods for reproducing the data analysis are available on GitHub.

### Code availability

ddHodge is implemented as a Julia package and is available at https://github.com/kazumits/ddHodge.jl. The Jupyter notebooks used to reproduce the figures and the related processed data in this manuscript are also available at https://github.com/kazumits/ddHodge_figures.

## Acknowledgments

We thank Profs. Yusuke Imoto and Yasuaki Hiraoka at Kyoto University for insightful discussions, and Advanced Research Initiative, Research Promotion Unit, Medical Institute of Bioregulation, Kyushu University, for their support. This work was supported by JST PRESTO JPMJPR2026, JST CREST JPMJCR23N3, JPMJCR24Q1, JSPS KAKENHI JP22H04696, JP23H04288 and JP25H01484 to K.M., AMED BINDS JP22ama121017j0001, JSPS KAKENHI JP24H02323, JP23H00372, JP22H04676 and JP22K19275 to Y.O. This work was also supported in part by the MEXT Promotion of Development of a Joint Usage / Research System Project: Pan-Omics DDRIC, MRCI for High Depth Omics, CURE:JPMXP1323015486 for MIB and RIIT in Kyushu University.

## Author contributions

K. M. and Y.O. conceived and interpretation of results, and K.M. built the theoretical framework of ddHodge and developed software, K. M. and Y.O. wrote the paper.

## Supplemental information

Figures S1–S3, and Notes of expository text of the mathematics in ddHodge.

