## Supplemental Information for "Geometry-preserving vector field reconstruction of high-dimensional cell-state dynamics using ddHodge"

#### 1 Supplemental Figures

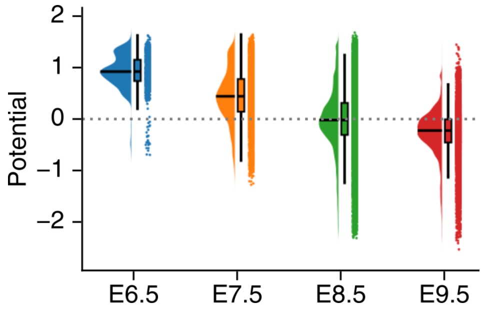

**Supplemental Figure S1**: Developmental stage and potential as shown in Figure 4D.

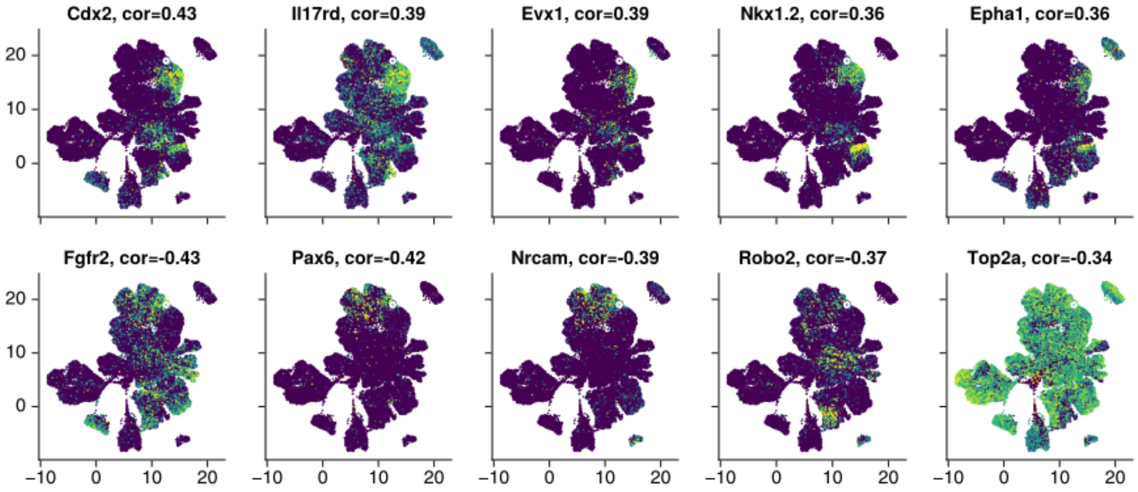

**Supplemental Figure S2**: Correlation between gene expression level and divergence in cluster 18. Each gene expression was shown as color. The correlateion coefficient are shown with each gene symbol as "[gene name], cor.=[value]". The marker of white circle indicate the location of POI.

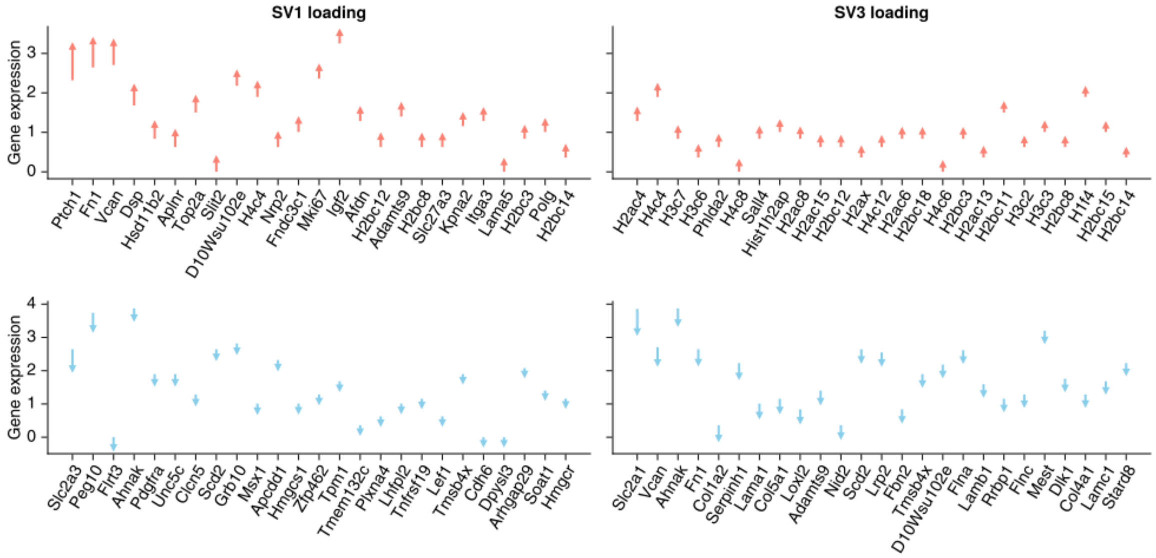

**Supplemental Figure S3**: Loadings of Schur vectors of the POI in cluster 14. The arrows represents direction and the length represents magnitude of infinitesimal change at POI represented by Schur vectors. The Y-axis indicates the gene expression level at POI. Each top 25 genes of positive and negative weights were shown here.

### 2 Supplemental Notes

Details and graphical explanations of the ddHodge method are provided in the following notes.

#### 2.1 Combinatorial Hodge decomposition

Hodge decomposition is a mathematical theorem that states that a vector field $\omega$ on a manifold $\mathcal{M}$ can be decomposed into three orthogonal components: a gradient grad $\alpha$, a divergence-free rotational component $\mathrm{curl}^*\beta$, and a residual, known as the harmonic component $\gamma$. Although the Hodge theory was built abstractly in terms of differential forms, the framework can be made computer friendly by discretizing the differential forms as a collection of the numerical values of the elements of a mesh or graph.

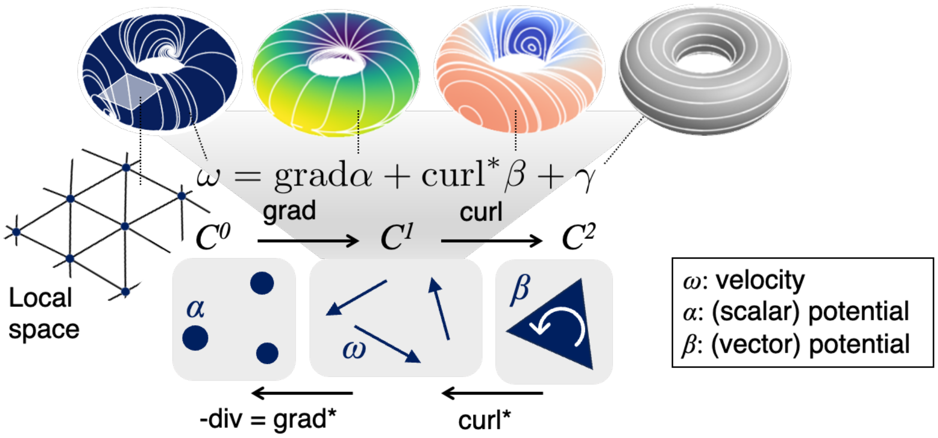

In particular, the estimation of the potential $\boldsymbol{u} \in \mathbb{R}^N$ corresponding to the scalar field $\alpha$ that gives the gradient component of the vector field requires only the construction of an $M \times N$ matrix $\delta_0 = \mathrm{grad}$ called the gradient operator from a given directed graph $\mathcal{G} = (V, E)$. Once we have $\delta_0$ and the data of the edge weights $\boldsymbol{w} \in \mathbb{R}^M$, we can obtain $\boldsymbol{u}$ via the least-squares method [13-14]:

$$\min_{\boldsymbol{u}} \|\delta_0 \boldsymbol{u} - \boldsymbol{w}\|^2$$

where $N$ is the number of vertices and $M$ is the number of edges in graph $\mathcal{G}$. This method, known as HodgeRank, was proposed for ranking problems and can also be viewed as a least squares solution to the discretized Poisson equation $\Delta f = \mathrm{div} g$.

In combinatorial Hodge decomposition, cochains $C^0, C^1, C^2$ are defined as spaces of functions used to assign real values to vertices (0-simplices), edges (1-simplices), and triangles (2-simplices), respectively.

The gradient operator $\delta_0 : C^0 \to C^1$ is a linear coboundary operator that can be represented as a matrix $\delta_0 \in \mathbb{R}^{M \times N}$ by considering $C^0 = \mathbb{R}^N$ and $C^1 = \mathbb{R}^M$. Specifically, for an edge $e = (i, j) \in E$, the matrix $\delta_0$ is constructed using elements $\{-1, 0, 1\}$ such that $(\delta_0 \boldsymbol{u})_e = u_j - u_i$. Although $\mathcal{G}$ is assumed to be a directed graph, this approach can also be applied to an undirected graph by adopting a consistent rule for the edge orientation, such as by regarding $(i, j)$ as a directed edge when $i < j$. Additionally, the alternativity $w_{(i,j)} = -w_{(j,i)}$ must be satisfied for the edge weights. Similarly, the curl operator $\mathrm{curl} = \delta_1 : C^1 \to C^2$, which takes the sum of the weights of the edges in the counterclockwise direction, can be represented by a matrix $\delta_1 \in \mathbb{R}^{L\times M}$ with elements $\{-1, 0, 1\}$.

In addition to the coboundary operators, for $x \in C^{k-1}$ and $y \in C^k$ with $k \in \{1, 2\}$, the adjoint of the coboundary operators is defined such that the inner product $\langle \delta_k x, y\rangle = \langle x, \delta_k^* y\rangle$ is preserved, resulting in the generation of a mapping with the reverse direction: $\delta_k^* = C^k \to C^{k-1}$. In the framework of combinatorial Hodge decomposition, these adjoints can be simply constructed as a matrix transpose $\delta_k^* = \delta_k^\top$. The divergence div $= -\delta_0^\top$ is the sum of the weights of the edges incident to the vertex, analogous to the usual definition of the divergence as the total flux per unit volume. Note that even if $\mathcal{G}$ is constructed using points in Euclidean space, the volume in the original space is not considered in the combinatorial Hodge decomposition.

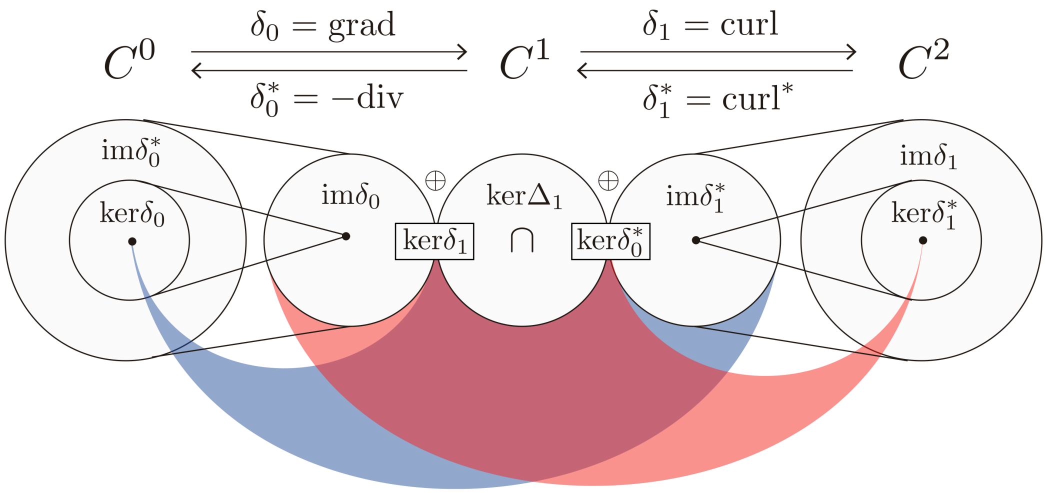

On the basis of these definitions of cochains, coboundary operators, and their adjoints, the property $\delta_k\delta_{k-1} = \delta_{k-1}^*\delta_k^* = 0$ always holds [1], allowing the vector space $C^1 = \mathbb{R}^M$ to be decomposed into the direct sum of the image of $\delta_0$ (gradient component), the image of $\delta_1^*$ (curl component), and the kernel of $\Delta_1$ (harmonic component), thus providing the same mathematical structure as Hodge decomposition for functions on manifold. Here, $\Delta_k = \delta_{k-1}\delta_{k-1}^* + \delta_k^*\delta_k$ is the abstract Hodge $k$-Laplacian [2].

The combinatorial Hodge decomposition approach, which provides a good estimation method of the potential, can be applied to vector fields defined by differential equations in continuous space. However, since the divergence and curl are the quantities per unit volume/area, they cannot be directly used as is. On the other hand, with numerical methods, which are preferable approaches when dealing with differential equations, challenges in constructing meshes and evaluating volumes in high-dimensional spaces are often encountered. Therefore, establishing methods for estimating the divergence and curl of vector fields via Hodge decomposition on graphs while avoiding the difficulties of mesh construction and volume evaluation is a key challenge addressed by ddHodge.

[1] Similar to $dd = \delta\delta = 0$ in exterior calculus

[2] When $k = 0$, $\Delta_0 = \delta_0^*\delta_0$ is the graph Laplacian.

##### 2.2 Defining edge weights from a given velocity

We focus on estimating the potential function $U : \mathbb{R}^N \rightarrow \mathbb{R}$ from the given velocity $\boldsymbol{v}_i$ for each point $\boldsymbol{x}_i$ in a sample of $N$ points in $d$-dimensional vector space.

The edge weights $w_{(i,j)}$ are defined as follows:

$$w_{(i,j)} = \frac{1}{2} \left(\boldsymbol{v}_i + \boldsymbol{v}_j\right)^\top \left(\boldsymbol{x}_j - \boldsymbol{x}_i\right)$$

From the perspective of discretization, this definition approximates the integral of the flux along an edge from $\boldsymbol{x}_i$ to $\boldsymbol{x}_j$. Geometrically, this definition gives an approximation of the potential difference $U\left(\boldsymbol{x}_j\right) - U\left(\boldsymbol{x}_i\right)$.

Let $\boldsymbol{\eta} = \boldsymbol{x}_j - \boldsymbol{x}_i$ be the vector indicating the direction of the edge. The projection of $\boldsymbol{v}_i$ onto $\boldsymbol{\eta}$ is given by $\boldsymbol{v}_i^\top \boldsymbol{\eta} = \boldsymbol{v}_i^\top \frac{\boldsymbol{\eta}}{\|\boldsymbol{\eta}\|} \|\boldsymbol{\eta}\|$. If the velocity is determined solely by the gradient of $U$, i.e., $\boldsymbol{v}_i = -\nabla U\left(\boldsymbol{x}_i\right)$, this projection gives the coefficient of the directional derivative $\nabla_{\boldsymbol{\eta}} U\left(\boldsymbol{x}_i\right) = -\boldsymbol{v}_i^\top \boldsymbol{\eta} / \|\boldsymbol{\eta}\|$. Thus, $w_{(i,j)}$ represents the average slope of $U$ along $\boldsymbol{\eta}$, which is then multiplied by the length $\|\boldsymbol{\eta}\|$.

In other words, $w_{(i,j)}$, as the average slope, determines the increment in the height of $U$ along $\boldsymbol{\eta}$. The shape of the potential function, for which the second derivative is a constant, corresponds to a quadratic function.

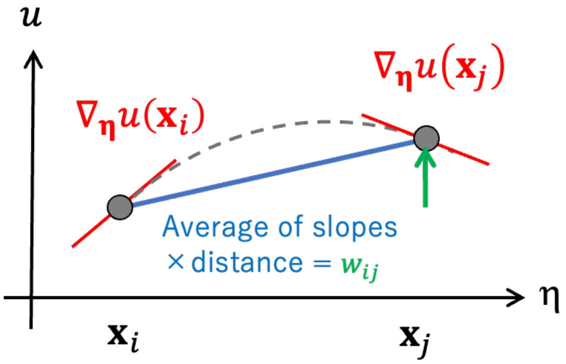

Consider a quadratic function along the axis of $\boldsymbol{\eta}$ with height $u(x)$ at position $x$:

$$u(x) = a(x - c)^2 + b$$
$$u'(x) = 2a(x - c)$$

The increment in the height $u\left(x_2\right) - u\left(x_1\right)$ can be obtained using the differential coefficients at each point $d_1 = u'\left(x_1\right)$ and $d_2 = u'\left(x_2\right)$:

$$u\left(x_2\right) - u\left(x_1\right) = a\left(x_2 - c\right)^2 - a\left(x_1 - c\right)^2 = \frac{d_2^2 - d_1^2}{4a}$$

Given $d_2 - d_1 = 2a\left(x_2 - x_1\right)$, substituting $a = \left(d_2 - d_1\right)/2\left(x_2 - x_1\right)$ yields the definition of the edge weight as follows:

$$u\left(x_2\right) - u\left(x_1\right) = \frac{1}{2}\left(d_1 + d_2\right)\left(x_2 - x_1\right)$$

The following figure visualizes the shape of the potential function $U$ (double-well potential), which is represented by piecewise quadratic functions fitted to the intervals of the sampled points and their slopes.

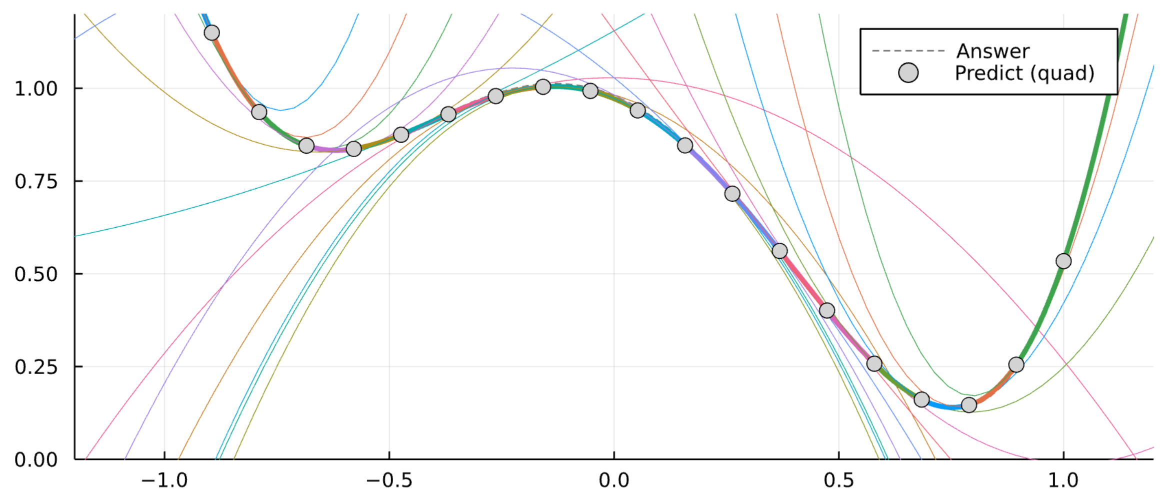

#### 2.3 Reconstruction of the Hessian matrix of $U$

The combinatorial divergence derived from the adjoint (transpose) of the gradient $\delta_0^\top$ differs from the usual divergence used in vector analysis. This is because the sum of the fluxes (weights) at the edges of the vertex does not match the definition of the divergence as the outgoing flux per unit volume. Leveraging the successful estimation of the potential using piecewise quadratic functions, we construct a div operator that can accurately restore the divergence in the original space.

From the previous results, the second-order derivative of the quadratic function can be easily derived as:

$$u''(x) = 2a = \frac{d_2 - d_1}{x_2 - x_1}$$

Since $x_1, x_2, d_1, d_2$ are all considered along the axis of $\boldsymbol{\eta}$, the second-order differential coefficient of the fitted quadratic function can be obtained by projecting the velocities onto $\boldsymbol{\eta}$ as follows:

$$U_{\eta\eta} = \frac{(\boldsymbol{x}_j - \boldsymbol{x}_i)^\top (\boldsymbol{v}_i - \boldsymbol{v}_j)}{\|\boldsymbol{x}_j - \boldsymbol{x}_i\|^2}$$

Note that the sign was flipped here: $d_2 - d_1 = \{(-\boldsymbol{v_j}) - (-\boldsymbol{v_i})\}^\top \boldsymbol{\eta}/\|\boldsymbol{\eta}\|$.

Since the divergence is calculated from the trace of the Hessian matrix $U_{xx} + U_{yy} + \cdots$, we next need to obtain the second-order partial derivatives $U_{xx}, U_{yy}$ in the original coordinate system $(x, y)$.

For simplicity, consider a linear relationship between the two-dimensional coordinate systems $(x, y)$ and $(\varphi, \psi)$. The relationship between the original Cartesian coordinates $(x, y)$

and an oblique coordinate system $(\varphi, \psi)$ spanned by two independent edges can be written as follows:

$$x = a\varphi + b\psi$$
$$y = c\varphi + d\psi$$

Next, consider the partial derivatives of $U\left(x(\varphi,\psi), y(\varphi,\psi)\right)$ with respect to $\varphi$ and $\psi$. Using the chain rule, we obtain:

$$\begin{aligned}\frac{\partial U}{\partial \varphi} &= \frac{\partial U}{\partial x}\frac{\partial x}{\partial \varphi} + \frac{\partial U}{\partial y}\frac{\partial y}{\partial \varphi}\\ &= a\frac{\partial U}{\partial x} + c\frac{\partial U}{\partial y}\end{aligned}$$

Similarly, for the second-order partial derivatives, we obtain:

$$\begin{aligned}\frac{\partial^2 U}{\partial \varphi^2} &= \frac{\partial}{\partial \varphi}\left(a\frac{\partial U}{\partial x} + c\frac{\partial U}{\partial y}\right)\\ &= a\frac{\partial}{\partial x}\frac{\partial U}{\partial \varphi} + c\frac{\partial}{\partial y}\frac{\partial U}{\partial \varphi}\\ &= \left(a\frac{\partial}{\partial x} + c\frac{\partial}{\partial y}\right)\left(a\frac{\partial U}{\partial x} + c\frac{\partial U}{\partial y}\right)\\ &= a^2\frac{\partial^2 U}{\partial x^2} + 2ac\frac{\partial^2 U}{\partial x \partial y} + c^2\frac{\partial^2 U}{\partial y^2}\end{aligned}$$

Focusing on the coefficients, the transformation of the second-order partial differential coefficients can be written in quadratic form. Thus, the relationship between the second-order directional derivative $U_{\eta\eta}$ with $\boldsymbol{\eta} = (a, c) = (\eta_1, \eta_2)$ and the Hessian matrix in the original coordinate system is given by:

$$\begin{aligned}U_{\eta\eta} &= (\eta_1\ \eta_2)\begin{pmatrix}U_{xx} & U_{xy}\\ U_{xy} & U_{xx}\end{pmatrix}\begin{pmatrix}\eta_1\\ \eta_2\end{pmatrix}\\ &= \eta_1^2 U_{xx} + 2\eta_1\eta_2 U_{xy} + \eta_2^2 U_{yy}\end{aligned}$$

By gathering the known second-order differential coefficients such as $U_{\eta\eta}$ in independent directions $\varphi, \psi, \ldots$, a linear equation to estimate the elements of $H$ can be obtained.

$$\begin{pmatrix}\varphi_1^2 & 2\varphi_1\varphi_2 & \varphi_2^2\\ \psi_1^2 & 2\psi_1\psi_2 & \psi_2^2\\ \eta_1^2 & 2\eta_1\eta_2 & \eta_2^2\end{pmatrix}\begin{pmatrix}U_{xx}\\ U_{xy}\\ U_{yy}\end{pmatrix} = \begin{pmatrix}U_{\varphi\varphi}\\ U_{\psi\psi}\\ U_{\eta\eta}\end{pmatrix}$$

We denote the equation as $C\boldsymbol{\beta} = \boldsymbol{y}$. The divergence is obtained from $-\mathrm{tr}H$ after estimating $H$.

In practice, the problem is better solved as a least square problem $\min_{\boldsymbol{\beta}} \|C\boldsymbol{\beta} - \boldsymbol{y}\|^2$ to mitigate issues in real data analysis, such as noise and limited sample size.

In conclusion, we obtain a statistical estimation of the Hessian matrix $H$ of $U$ without directly addressing the mesh design, area/volume calculation, or boundary conditions, relying solely on the sampling of the second-order partial differential coefficients from quadratic functions fitted along incident edges.

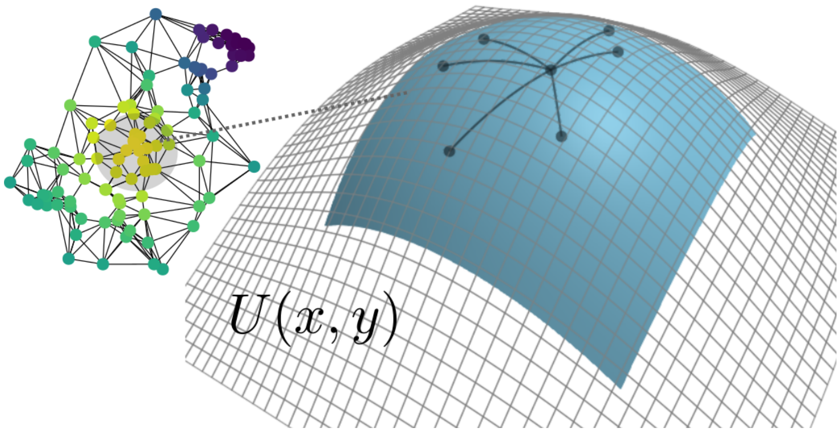

##### 2.4 $l_2$ regularization on a graph

To enable the regularization of the estimated values (Hessian matrix elements) between neighboring vertices, we merged the individual Hessian matrix estimation problems into a single linear equation:

$$C\boldsymbol{\beta} = \begin{pmatrix} C_1 & & \\ & \ddots & \\ & & C_N \end{pmatrix} \begin{pmatrix} \boldsymbol{\beta}_1 \\ \vdots \\ \boldsymbol{\beta}_N \end{pmatrix}$$

This allows us to introduce a penalty (Dirichlet energy on the graph) to the squares of the differences $\boldsymbol{\beta}_i - \boldsymbol{\beta}_j$ between adjacent vertices as:

$$P(\boldsymbol{\beta}) = (D_0\boldsymbol{\beta})^\top (D_0\boldsymbol{\beta}) = \boldsymbol{\beta}^\top L_0 \boldsymbol{\beta}$$

Here, assume that the $p$ parameters are arranged in a certain order as $\boldsymbol{\beta}_i = \left(U^i_{xx}, U^i_{xy}, U^i_{yy}\right)$, such that the differences between each parameter can be individually evaluated using the extended gradient matrix $D_0 = \delta_0 \otimes I_p$ with the Kronecker product and $L_0 = D_0^\top D_0$.

The optimization problem with this regularization term is then solved as:

$$\min_{\boldsymbol{\beta}} \|\boldsymbol{y} - C\boldsymbol{\beta}\|_2^2 + \lambda P(\boldsymbol{\beta})$$

with some small constant $\lambda \in \mathbb{R}$.

The following solution can be obtained from the extrema with respect to the parameter $\boldsymbol{\beta}$:

$$\widehat{\boldsymbol{\beta}} = \left(C^\top C + \lambda L_0\right)^{-1} C^\top \boldsymbol{y}$$

Given the large $Np \times Np$ size of the sparse matrix $L_0$, efficient solvers such as the Krylov subspace method are recommended.

##### 2.5 Reconstruction of the curl

For differential forms in the Euclidean space $\mathbb{R}^2$, the relationship between div and curl is given by:

$$\mathrm{div}(\star\xi) = (\star d\star)(\star\xi) = (\star d)(\star\star)\xi = -\mathrm{curl}\xi$$

where $d$ is the exterior derivative and $\star$ is the Hodge star operator. Therefore, the curl can be approximated by the divergence of the *dual velocity field* $\star\xi$, which is obtained by rotating the velocity by $\pi/2$ in the PC1-2 plane.

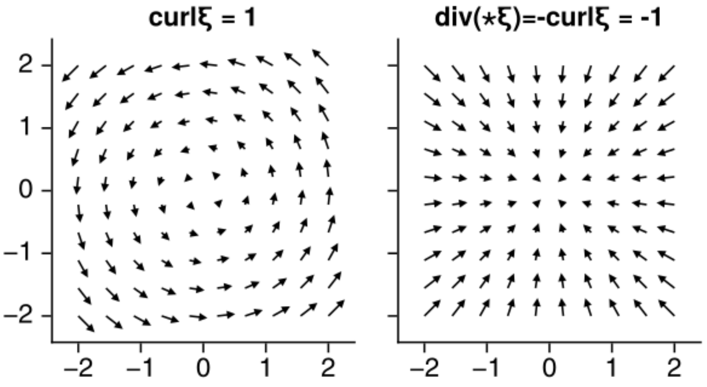

#### 2.6 Reconstruction of the Jacobian matrix

According to the result of Hodge decomposition, a vector field $f$ can be expressed as the sum of a gradient part $-\nabla U$ and a divergence-free residual part $r$:

$$f = -\nabla U + r$$

Here, let $D$ be the operator that gives the Jacobian matrix of $f$. Since $D$ is linear and $D\nabla U$ is the Hessian matrix $H$ of $U$, $D(f)$ has the following decomposition:

$$D(f) = D(-\nabla U + r) = -H + D(r)$$

If $r$ is in 2D Euclidean space and consists of only rotational components (i.e., there are no holes in the space), $D(r)$ can be expressed as:

$$D(r) = \frac{\theta}{2}Q$$

where $Q$ is the infinitesimal rotation matrix:

$$Q = \begin{pmatrix} 0 & -1 \\ 1 & 0 \end{pmatrix},$$

$$\exp(\theta Q) = \begin{pmatrix} \cos\theta & -sin\theta \\ \sin\theta & \cos\theta \end{pmatrix}$$

Here, $\theta \in R$ is the magnitude of the rotation, which is calculated from the previously derived *curl*. The factor 1/2 in $D(r)$ ensures that when $\theta = 1$, $\mathrm{curl}(r) = (Q_{21} - Q_{12})/2 = 1$.

At this time, we do not perform a complete reconstruction of the rotational components in dimensions higher than 3. Instead, we approximate the rotation only in the PC1-2 plane of each tangent space. To obtain the $m \times m$ matrix of $D(r)$, we use the basis transformation

$$Q' = (W^\top W)^{-1} W^\top S Q (S^\top S)^{-1} S^\top W$$

This transformation maps the rotation $Q \in SO(2)$ on the PC1-2 plane to the rotation $Q' \in SO(m)$ on the same plane in the $m$-dimensional tangent space. Here, $S$ is the matrix composed of the basis of the PC1-2 plane, and $W$ is the matrix composed of the basis of the $m$-dimensional tangent space.

In conclusion, the Jacobian matrix of $f$ in the $m$-dimensional tangent space can be approximated as follows:

$$D(f) \approx -H + \frac{\theta}{2} Q'$$

#### 2.7 Designing the parallel coordinate system

To perform $l_2$ regularization on the data manifold (a set of local PCA spaces), it is necessary to establish local coordinate systems that provide correspondence between adjacent tangent vector spaces. In other words, in a curved space, determining the direction in which smoothness is required is not straightforward. This issue is addressed by the concept of *parallel transport* in differential geometry.

In this section, we introduce the idea of near parallel coordinate design in the $n$-dimensional space by using the subspace alignment method employed with vector diffusion maps [19] and the approach for designing discrete connection Laplacians via sheaves on graphs [25-26], which is an extension of combinatorial Hodge decomposition. The procedure consists of two steps: (1) subspace alignment and (2) minimization of the Rayleigh quotient of the discrete connection Laplacian.

##### Step 1: Subspace alignment

For two vertices $i$ and $j$ in a graph, let $O_i$ and $O_j$ be the $d \times r$ matrices composed of the orthonormal basis of the $r$-dimensional tangent vector spaces $T_i\mathcal{M}$ and $T_j\mathcal{M}$, respectively. The vector spaces spanned by the columns of $O_i$ and $O_j$ are denoted as $Col\,(O_i) = T_i\mathcal{M}$ and $Col\,(O_j) = T_j\mathcal{M}$.

If the two spaces have a flat connection, i.e., $Col\,(O_i) = Col\,(O_j)$, then there exists an orthogonal matrix $O_{ij}$ that maps $O_i$ into $O_j$ via the linear combination of the columns of $O_i$:

$$\begin{aligned} O_i O_{ij} &= O_j \\ O_{ij} &= O_i^\top O_j \end{aligned}$$

Thus, for any vector $\boldsymbol{z_i} \in Col\,(O_i)$, $O_{ij}$, called the *transfer matrix,* provides the alignment of the axes in different spaces $\boldsymbol{z_j} = {O_{ij}}^\top \boldsymbol{z}_i \in Col\,(O_j)$ such that $O_i \boldsymbol{z}_i = O_j \boldsymbol{z}_j$.

However, in general, $Col\,(O_i) \neq Col\,(O_j)$ when the space has a nonzero curvature, and a linear combination cannot provide such alignment. Therefore, an approximate version of the transfer matrix is obtained as follows:

$$\tilde{O}_{ij} = \underset{O \in O(r)}{\operatorname{argmin}} \left\| O - O_i^\top O_j \right\|_F^2$$

where $\| \cdot \|_F$ is the Frobenius norm and $O(r)$ is the orthogonal group. The solution is obtained via singular value decomposition (SVD):

$$O_i^\top O_j = U \Sigma V^\top$$
$$\tilde{O}_{ij} = U V^\top = \tilde{O}_i^\top \tilde{O}_j$$

where $\tilde{O}_i = U^\top$ and $\tilde{O}_j = V^\top$.

The columns of $U$ and $V$ corresponding to the largest singular value $\sigma$ provide the pair $\boldsymbol{u}$ and $\boldsymbol{v}$ that maximizes $\cos\theta = \langle O_i \boldsymbol{u}, O_j \boldsymbol{v} \rangle = \sigma \leq 1$, where $\theta$ is the angle between $O_i \boldsymbol{u}$ and $O_j \boldsymbol{v}$. This is equivalent to canonical correlation analysis (CCA), which is used to find the closest axis pair in the column spaces of the given matrices $X$ and $Y$. Additionally, the angles $\theta_i = \operatorname{acos}(\sigma_i)$ corresponding to the singular values $\sigma_i$ are called canonical angles, and the magnitude $\|\boldsymbol{\theta}\| = \|(\theta_i, \ldots, \theta_r)\|$ can be used as the **Grassmann distance** to measure the distance between different subspaces.

##### Step 2: Minimizing the Rayleigh quotient of the connection Laplacian

The above method provides parallel transport between a pair of tangent spaces. However, since the solution depends on the selection of the pair (cf. holonomy), it cannot be used as a consistent local coordinate system.

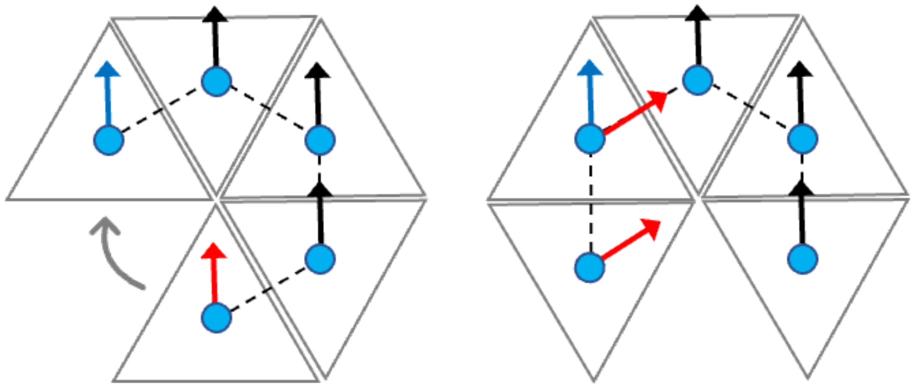

The above figure illustrates the procedure of parallel transport on a developed surface of a 3D cone approximated by five tangent triangle planes. The observed holonomy (inconsistency after traverse) is caused by the angular defect.

Instead, we consider obtaining pairs of $\boldsymbol{z}_i \in Col(O_i)$ and $\boldsymbol{z}_j \in Col(O_j)$ for all edges $(i, j) \in E$ that are as close to parallel transport as possible. Let the *deviation from parallel transport* be $\boldsymbol{\varepsilon}_{(i,j)} = \tilde{O}_i \boldsymbol{z}_i - \tilde{O}_j \boldsymbol{z}_j$.

We aim to minimize:

$$\|\boldsymbol{\varepsilon}_{(i,j)}\|^2 = \left(\tilde{O}_i z_i - \tilde{O}_j \boldsymbol{z}_j\right)^\top \left(\tilde{O}_i \boldsymbol{z}_i - \tilde{O}_j \boldsymbol{z}_j\right)$$
$$= 2\left(1 - \boldsymbol{z}_i^\top \tilde{O}_i^\top \tilde{O}_j \boldsymbol{z}_j\right)$$

where $\|\boldsymbol{z}_i\|^2 = \|\boldsymbol{z}_j\|^2 = 1$. This becomes the problem of maximizing $\boldsymbol{z}_i^\top \tilde{O}_i^\top \tilde{O}_j \boldsymbol{z} j$. According to the previous results, if we select $\boldsymbol{z}_i = \boldsymbol{u}$ and $\boldsymbol{z}_j = \boldsymbol{v}$ via SVD, where $\boldsymbol{z}_i^\top \tilde{O}_i^\top \tilde{O}_j \boldsymbol{z}_j = 1$, we can still check that perfect parallel transport is possible with $\|\boldsymbol{\varepsilon}(i,j)\|^2 = 0$.

If there is only one pair of tangent spaces, we can select $\boldsymbol{u}$ and $\boldsymbol{v}$ as described. However, if vertex $j$ also has an edge $(k,j)$, different solutions $\boldsymbol{v'} \neq \boldsymbol{v}$ are given by the SVD of $O_k^\top O_j$. Keeping $\boldsymbol{z}_j = \boldsymbol{v} \neq \boldsymbol{v'}$ may introduce a large deviation $\|\boldsymbol{\varepsilon}(j,k)\|^2$. To prevent this, we need to determine $\boldsymbol{z}_i, \boldsymbol{z}_j, \boldsymbol{z}_k, \ldots \in \mathbb{R}^r$ that minimize the total deviation $\sum_{e\in E} \|\boldsymbol{\varepsilon}_e\|^2$ over all edges. This type of optimization problem on a graph with vector-valued functions $\boldsymbol{z}_i$ on vertices can be addressed using the framework of sheaves on graphs.

Similar to Hodge decomposition, $\varepsilon_e$ can be evaluated on the basis of the sheaf version of the gradient $\delta_s$ of the selected coordinates $\boldsymbol{z}_i$ as follows:

$$\delta_s \boldsymbol{z} = \begin{pmatrix} * & \cdots & & & \\ O & \mathcal{F}_{i\trianglelefteq e} & O & -\mathcal{F}_{j\trianglelefteq e} & O \\ & & & \cdots & * \end{pmatrix} \begin{pmatrix} * \\ \boldsymbol{z}_i \\ * \\ \boldsymbol{z}_j \\ * \end{pmatrix} = \begin{pmatrix} * \\ \mathcal{F}_{i\trianglelefteq e}\boldsymbol{z}_i - \mathcal{F}_{j\trianglelefteq e}\boldsymbol{z}_j \\ * \end{pmatrix}$$

In combinatorial Hodge decomposition, the gradient $\delta_0$ represents the difference $z_j - z_i \in \mathbb{R}$ between vertices $i$ and $j$ for edge $(i,j)$. In contrast, this gradient $\delta_s$ generalizes $\delta_0$ to a block matrix form, allowing the evaluation of differences $\mathcal{F}_{i\trianglelefteq e}\boldsymbol{z}_i - \mathcal{F}_{j\trianglelefteq e}\boldsymbol{z}_j$ with vector-valued functions $\boldsymbol{z_i}, \boldsymbol{z_j} \in \mathbb{R}^r$ at the vertices. These linear transformations $\mathcal{F}_{i\trianglelefteq e}$ and $\mathcal{F}_{j\trianglelefteq e}$ are called restriction maps.

Thus, by setting $\mathcal{F}_{i\trianglelefteq e} = \tilde{O}_i$ and $\mathcal{F}_{j\trianglelefteq e} = \tilde{O}_j$ for each row of $\delta_s$, we can concisely describe the total deviation over the graph as:

$$\mathrm{loss}\,(\boldsymbol{z}) = \sum_{(i,j)\in E} \|\tilde{O}_i \boldsymbol{z}_i - \tilde{O}_j \boldsymbol{z}_j\|^2 = \boldsymbol{z}^\top \Delta_s \boldsymbol{z}$$

where $\Delta_s = \delta_s^\top \delta_s$ is called the sheaf Laplacian.

The minimization of $\mathrm{loss}(\boldsymbol{z})$ while keeping the magnitude of $\boldsymbol{x}$ constant is known as Rayleigh quotient $\left(\boldsymbol{z}^\top \Delta_s \boldsymbol{z}\right) / \left(\boldsymbol{z}^\top \boldsymbol{z}\right)$ minimization. By definition, $\Delta_s$ is positive semidefinite, so it always has a nonnegative minimum eigenvalue, and the minimum value of the Rayleigh quotient is the smallest eigenvalue of $\Delta_s$.

In conclusion, we adopted the eigenvectors $\boldsymbol{z}_i$ corresponding to the smallest eigenvalues of $\Delta_s$ as the local coordinate axes, which are nearly parallel and consistent across all tangent spaces.
